# Modeling the marmoset brain using embryonic stem cell-derived cerebral assembloids

**DOI:** 10.1101/2023.02.28.530008

**Authors:** Tomoki Kodera, Ryosuke F. Takeuchi, Sara Takahashi, Keiichiro Suzuki, Hidetoshi Kassai, Atsu Aiba, Seiji Shiozawa, Hideyuki Okano, Fumitaka Osakada

## Abstract

Studying the non-human primate (NHP) brain is required for the translation of rodent research to humans, but remains a challenge for molecular, cellular, and circuit-level analyses in the NHP brain due to the lack of *in vitro* NHP brain system. Here, we report an *in vitro* NHP cerebral model using marmoset (*Callithrix jacchus*) embryonic stem cell-derived cerebral assembloids (CAs) that recapitulate inhibitory neuron migration and cortical network activity. Cortical organoids (COs) and ganglionic eminence organoids (GEOs) were induced from cjESCs and fused to generate CAs. GEO cells expressing the inhibitory neuron marker LHX6 migrated toward the cortical side of CAs. COs developed their neural activity from a synchronized pattern to an unsynchronized pattern as COs matured. CAs showed mature neural activity with an unsynchronized pattern. The marmoset assembloid system will provide an *in vitro* platform for the NHP neurobiology and facilitate translation into humans in neuroscience research, regenerative medicine, and drug discovery.

## 1. Introduction

Clinical translation of regenerative medicine and drug development, as well as understandings of human brain function and disease, require biological analyses in nonhuman primates (NHPs) [1]. The drug development success rate at clinical trials in psychiatric disorders is <10% [2]. This low success rate could be attributed to differences in brain structure and function between rodents and humans. To improve this issue, both *in vivo* and *in vitro* models of not only rodents, but also NHPs are required for evaluating drug effects. Moreover, the clinical translation of transplantation therapies also needs NHP studies to evaluate their safety and effectiveness. In particular, autologous and allogeneic transplantation systems of differentiated cells or organoids derived from induced pluripotent stem cells (iPSCs) using NHPs are crucial for regenerative medicine development [3]. Recently, great attention has been directed to the common marmoset (*Callithrix jacchus)* as a powerful animal model that bridges mice and humans for biomedical and neuroscience research applications [4]. Marmosets share significant brain structure and function features with humans, such as the prefrontal cortex, which is involved in higher cognitive function [5]. The lack of *in vitro* marmoset models for molecular and cellular analyses has limited marmoset research development, despite the recent increased attention to pathophysiological and behavioral studies in marmosets, as well as transgenic marmoset generation [6,7]. Therefore, *in vitro* generation of differentiated neurons and brain organoids from marmosets, if possible, would contribute to neuroscience research, drug development, and regenerative medicine.

The mammalian cerebral cortex is responsible for higher-order brain functions, including sensory perception, motor control, and cognition, which are generated by computations of excitatory and inhibitory neurons [8,9]. During early mammalian development, excitatory glutamatergic neurons are generated in the dorsal telencephalon, while inhibitory GABAergic neurons are generated in the ventral telencephalon [10–12]. GABAergic neurons tangentially migrate toward the dorsal telencephalon and integrate into cortical circuits. Monkey and human cerebral cortex are highly expanded compared with other mammals, such as mice [13]. Human brain development and function features implicated in multiple neurodevelopmental disorders are not similar to rodent models [14,15]. Many attempts to analyze these developmental processes, cortical circuit organization and function, and circuit malfunction in humans have been made using human embryonic stem cells (ESCs) and iPSCs because of the invasiveness to human bodies and human tissue inaccessibility [16–18]. Organoids are powerful in vitro models for the three-dimensional brain [19]. Recently, Biery et al., and Xiang et al. have demonstrated that assembled human organoids by cortical organoid (CO) and subpallium organoid fusion, called assembloids, mimic the interneuron migration from the subpallium organoid to the CO *in vitro* [20–22]. Although human ESCs and iPSCs have been differentiated into cortical neurons in two-dimensional cultures and organoids in three-dimensional cultures, *in vitro* functional cortical circuit construction remains challenging.

In this study, we developed a marmoset brain *in vitro* system based on marmoset ESC (cjESC)-derived organoids to model the excitatory and inhibitory neuron interaction in the marmoset cerebral cortex, a pivotal event of cortical development and function. COs and ganglionic eminence organoids (GEOs) derived from cjESCs were assembled to generate the cerebral assembloids (CAs) and reconstruct inhibitory neuron migration to the cortical structure. Moreover, COs change their neural activity from a synchronized pattern to an unsynchronized pattern as they mature, indicating COs exhibit developmental-stage-dependent changes in spontaneous neural activity. CAs exhibited mature neural activity with an unsynchronized pattern. Thus, marmoset CAs will offer a powerful *in vitro* platform to study NHP neurobiology, which supplements *in vivo* NHP work and presents translational benefits for neuroscience research, regenerative medicine, and drug development.

## 2. Materials and methods

### 2.1. cjESC culture

cjESCs (CMES40, RIKEN BRC) were maintained on a feeder layer of mitomycin C-treated mouse embryonic fibroblasts (MEFs). Mitomycin C-treated MEFs were prepared as described previously [23]. cjESCs were cultured with knockout DMEM (Thermo) containing 20% (v/v) knockout serum replacement (KSR; Thermo), 1% (v/v) MEM-NEAA (Wako), 1 mM L-glutamine (Wako), 100 U/ml penicillin, 100 μg/ml streptomycin (Wako), 0.2 mM 2-mercaptoethanol (Wako), and 10 ng/ml bFGF (Peprotech) in a humidified atmosphere of 5% CO_2_ at 37°C. cjESCs were passaged with 0.25% trypsin-EDTA (Wako) every 5 days.

### 2.2. Differentiation of cjES cells

On organoid culture day 0, cjESCs were dissociated to single cells in 0.25% trypsin-EDTA and reaggregated in a differentiation medium (10,000 cells/100 μl for each well) according to the SFEBq culture with some modifications [16]. The differentiation medium contained GMEM (Wako) and 20% (v/v) KSR supplemented with 1 mM pyruvate (Wako), 1% (v/v) MEM-NEAA, 100 U/ml penicillin, and 100 μg/ml streptomycin. For cortical organoid induction, 50 μM Y-27632 (medchemexpress) was added in the differentiation medium from days 0–3, 5 ng/ml bFGF from days 3–18, and 500 nM A 83 01 (Wako), 1 μM LDN193189 (Stemgent), and 3 μM IWR-1-*endo* (Wako) from days 0–18. For MGE organoid induction, 500 nM SAG (Santa cruz) was added from days 9–18 on top of the cortical organoid condition (Y-27632, bFGF, A-83-01, LDN193189, and IWR-1-*endo*). On day 18, aggregates were transferred to a low-binding 60-mm dish and cultured in a maturation medium, which contained a 1:1 mixture of DMEM/F12 (Sigma) and neurobasal (Thermo) supplemented with 0.5% N2 supplement (Wako) with Transferrin, 1% B27 supplement (Thermo), 0.5% nonessential amino acids, 2 mM L-glutamine, 100 U/ml penicillin, 100 μg/ml streptomycin, 2.5 μg/ml Insulin (Wako), 50 μM 2-mercaptoethanol, 200 mM ascorbic acid (Wako), and 20 ng/ml BDNF (Peprotech). From day 18, organoids were maintained on a bio orbital shaker at 65 rpm in a 5% CO_2_ and 40% O_2_ at 37°C humidified atmosphere.

### 2.3. Generation of cerebral assembloids

A CO and a GEO were transferred to a low-binding 1.5-ml tube containing the maturation medium added 1% (v/v) Matrigel (Corning) and placed in contact with each other for 4 days in an incubator of a 5% CO_2_ at 37°C humidified atmosphere. The formed assembloid was transferred to a low-binding 60-mm dish on an orbital shaker at 65 rpm and maintained with medium changes every 3 days.

### 2.4. Real-time quantitative PCR

Total organoids RNA was isolated using FavorPrep Tissue Total RNA Purification Mini Kit [24], and 500 ng total RNA was reverse transcribed using PrimeScript RT Master Mix (Takara). Synthesized cDNA was subjected to quantitative PCR using TB Green Fast qPCR Mix (Takara) on the LightCycler system (Roche). Primers used in the present study were listed in Supplementary Table 1.

### 2.4. Immunohistochemistry

Organoids were fixed in 4% paraformaldehyde at 4°C overnight and washed three times with PBS at room temperature. Fixed organoids were transferred to 15% sucrose for overnight incubation, followed by 30% sucrose until organoids were sunk in sucrose solution. Organoids were transferred to cryomold (Sakura Finetek) and embedded in a 2:1 mixture of O.C.T compound (Sakura Finetek) and 30% sucrose on dry ice. The organoids were sectioned at 18 μm on a cryostat (CM3050S, Leica). For immunostaining, the slices were reacted with Blocking One (Nacalai tesque) for 1 h at room temperature and then with primary antibodies overnight at 4°C. Slices were washed with PBS three times and incubated with secondary antibodies for 2 h at room temperature. Primary antibodies and their working dilutions were as follows: anti-SOX2 (1:1000, rabbit; Abcam), anti-ß-III TUBULIN (1:300, mouse; Abcam), anti-FOXG1 (1:500, rabbit; Takara), anti-MAP2 (1:500, chicken; Abcam), anti-TBR1 (1:400, rabbit; Cell signaling), anti CTIP2 (1:500, rat; Abcam), anti-REELIN (1:500, mouse; Abcam), anti-SATB2 (1:500, mouse; Abcam), anti-TBR2 (1:500, rabbit; Abcam), anti-GSH2 (1:200, rabbit; Abcam), anti-NKX2.1 (1:500, mouse; Santa Cruz), anti-LHX6 (1:100, mouse; Santa Cruz), anti-OLIG2 (1:500, rabbit; Abcam), anti-GAD67 (1:200, mouse; Millipore), anti-PV (1:500, Goat; Swant), anti-SST (1:150, rat; Millipore), anti-SYN1 (1:500, rabbit; Cell sigmaling), anti-PSD95 (1:300, mouse; Sigma), anti-GEPHYRIN (1:100, mouse; Synaptic systems), anti-GFP (1:1000, rabbit; Abcam; 1:1000, chicken; Abcam), anti-mCherry (1:1000, rat; Invitrogen), and anti-DsRed (1:1000, rabbit; Clontech). Secondary antibodies used were as follows: anti-mouse IgG, anti-rabbit IgG, anti-goat IgG, anti-rat IgG, anti-chicken IgY antibodies conjugated with Alexa Fluor 488, Alexa Fluor 594, or Alexa Fluor 647 (1:1000, donkey; Jackson). Cell nuclei were counterstained with DAPI. Specimens were imaged with confocal microscope (LSM800, Zeiss).

### 2.5. AAV production

AAV vectors were generated in HEK293T cells as described previously [25]. Briefly, AAV9 was produced by HEK293T cells transfected with pHelper, the AAV9 rep/cap vector, and the genomic vector pAAV-Esyn-tTA or pAAV-tetO-jGCaMP8m. Virus-producing cells were collected 3 days after transfection and lysed by freeze-and-thaw cycles for purification. After centrifugation, the supernatant was loaded onto gradients (15%, 25%, 40%, and 58%) of iodixanol OptiPrep (Serumwerk Bernburg). After centrifugation of 16,000 × *g* at 4°C for 4 h, 150–200 μl of the 40% iodixanol fraction was collected and used for infection experiments. AAVs genomic titers were quantified using quantitative PCR. AAV9-Esyn-tTA and AAV9-tetO-jGCaMP8m titers were 7.41 × 10^14^ and 3.31 × 10^14^ viral genome/ml, respectively.

### 2.6. Lentivirus production

HIV-based lentivirus (LV) was generated in HEK293T cells as described previously [26]. VSV-G-LV-CAG-tdTomato was produced by HEK293T cells transfected with the helper plasmids pMDL and pREV, the envelope expression plasmid pCMV-VSV-G, and the genomic vector pBOB-CAG-tdTomato. The supernatant from virus-producing cells was collected and filtered 40 h after transfection. After ultracentrifugation of 21,000 × *g* at 4°C for 2h, the pellet was resuspended in 180 μl of HBSS. VSV-G-LV-CAG-tdTomato titer was 2.13 × 10^7^ unit/m.

### 2.7. Two-photon calcium imaging

Approximately 50-day-old or 70-day-old organoids were subjected to calcium imaging. Organoids were infected with AAV9-Esyn-tTA and AAV9-tetO-jGCaMP8m for 3 days. The organoids lower half were embedded in 4% low melting point agarose 6 days after infection and filled with oxygenated artificial cerebrospinal fluid consisting of 125 mM NaCl, 2.5 mM KCl, 1.25 mM NaH_2_PO_4_-H_2_O, 25 mM NaHCO_3_, 25 mM D-Glucose, 2 mM CaCl_2_, and 1 mM MgCl_2_. Imaging was performed using a two-photon fluorescent microscope equipped with GaAsP-type non-descanned detector and resonant scanner (A1R-MP^+^, Nikon) [27]. A 920 nm pulse laser (InSight DeepSee+, Spectra-Physics) was used to excite jGCaMP8m. Time-lapse images with a 25× water immersion lens (NA: 1.1, Nikon) at a 512 × 512-pixel resolution were acquired at 30.0 fps for 90 s.

MATLAB (Mathworks, R2022b) was used to perform all image processing and statistical quantification. In the first image-processing step of time-lapse data, cross correlation based rigid image registration [28] was performed to correct the image displacement caused by motion artifacts. Fluorescent intensity fluctuation from individual cells was quantified. Regions-of-interest of individual cells were semi manually defined with the “Cell Magic Wand” plugin in ImageJ [29]. Fluorescence data were smoothed by sliding-moving-average (filter width = 5 frames) to reduce high-frequency artifact effect caused by electrical noise. To evaluate each neuron’s activity, normalized fluorescent change (dF/F0) was computed. Each cell baseline (F0) was computed by a low-pass percentile filter (10 percentile, cut-off frequency = 1/15 Hz). We recorded 1074 cells from six organoids (16 sites from 6 organoids) and defined 779 cells as “Active cells” for this study according to the following three criteria (Fig. 4G): First, ΔF/F distribution skewness during recording had to be >0.25. Second, signal-noise ratio had to be >0.001. Third, at least one calcium event was detected during the recording period by the deconvolution algorithm as below. The signal-noise ratio and calcium events were quantified with the OASIS algorithm in the CaImAn-MATLAB package [30]. Among the detected deconvolved events, events with 0.5 seconds or less interval were quantified as a single event. Activity sparseness index of neuron i is given by:

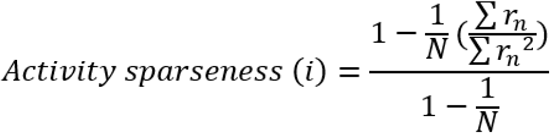

where, r_n_ is the ΔF/F for each time point, and N is the total number of time points. A neuron will have high lifetime sparseness if it is silent most of the time, but occasionally produces strong responses at only small subsets of time [31]. Pairwise correlation values for each site were quantified by the average of correlation coefficient between all active-cell pairs. Population coupling was quantified to characterize how different neurons relate to large-scale firing patterns. The population coupling for individual neurons was quantified by calculating Pearson’s correlation coefficient between individual neurons and all other simultaneously recorded neurons’ summed activity [32]. Principal component analysis was performed to assess how diverse the population activity patterns of neurons were.

### 2.8. Tissue clearing

For whole assembloid analyses, CUBIC protocol [33] was applied, followed by immunohistochemistry. Briefly, day 37-58 assembloids were fixed with 4% PFA at 4°C overnight. The next day, assembloids were washed with PBS three times and incubated with CUBIC Reagent-1 diluted 2-fold with H_2_O for 3 h and then with Reagent-1 overnight at 37°C. Assembloids were washed with PBS three times for 2 h and then stained with anti-GFP (1:1000) and anti-mCherry (1:1000) antibodies in PBS containing 0.1% Triton X-100 and 5% Blocking One for 2 days at 37°C. Assembloids were subsequently washed with PBS three times for 2 h and incubated with secondary antibodies for 2 days at 37°C. For refractive index matching, assembloids were incubated with 2-fold diluted CUBIC Reagent-2 for 3 h at room temperature and then with Reagent-2 overnight at room temperature.

### 2.9. Genome editing in cjESCs

CRISPR-Cas9 was used to generate cjESC reporter lines: AAVS1-EYFP-cjESCs and AAVS1-mCherry-cjESCs. Guide RNA (gRNA) expression vectors were constructed by assembling AflII-digested pCAG-mCherry-gRNA (Addgene, plasmid 87110) with gRNA sequences using NEBuilder HiFi DNA Assembly (NEB). Donor DNA plasmids were constructed by assembling BamHI and HindIII-digested pUC19 vector with left and right homology arm (HA) fragments, a CAG promoter fragment, and a fluorescent protein (EYFP or mCherry) fragment. Left and right HAs were amplified with PCR from the marmoset genome extracted from cjES cells. Other fragments were obtained by PCR amplification from appropriate plasmids and cjES cells (1.0 × 10^6^ cells, 100-mm dish: CMES40) were transfected with 6.25-μg of gRNA expression vector, 6.25 μg of Cas9 expression vector (pCAG-1BPNLS-Cas9-1BPNLS-2A-GFP; Addgene, plasmid 87109) and 12.5 μg of donor plasmid using Lipofectamine LTX (Thermo) according to the manufacturer’s protocol. The cjESCs were dissociated into single cells 2 days after transfection and replated on a 100 mm dish at 1.0 × 10^4^ cells. Around 2 weeks after replating, single cjESC colonies were manually picked up based on EYFP or mCherry expression and genotyped by PCR with primers listed in Supplementary Table 1. Knocked-in cjESCs subsequently performed expanded-culture and made stocks.

### 2.10. Statistical analysis

Values are expressed as means ± S.E.M. Statistical analyses were performed with Prism or MATLAB. Statistical significance of the difference between groups was determined by one-way ANOVA followed by Dunnett’s test (Fig. 1D) or Tukey’s test (Fig. 3E, 3F). Unpaired *t*-test was used to evaluate data from Fig. 2C, 2E, 2F, 3I, S1B, S1C, and S1E. Wilcoxon rank sum test was used to evaluate data from Fig. 4G-K and 5D-H. Probability values <5% were considered significant.

**Fig. 1:**
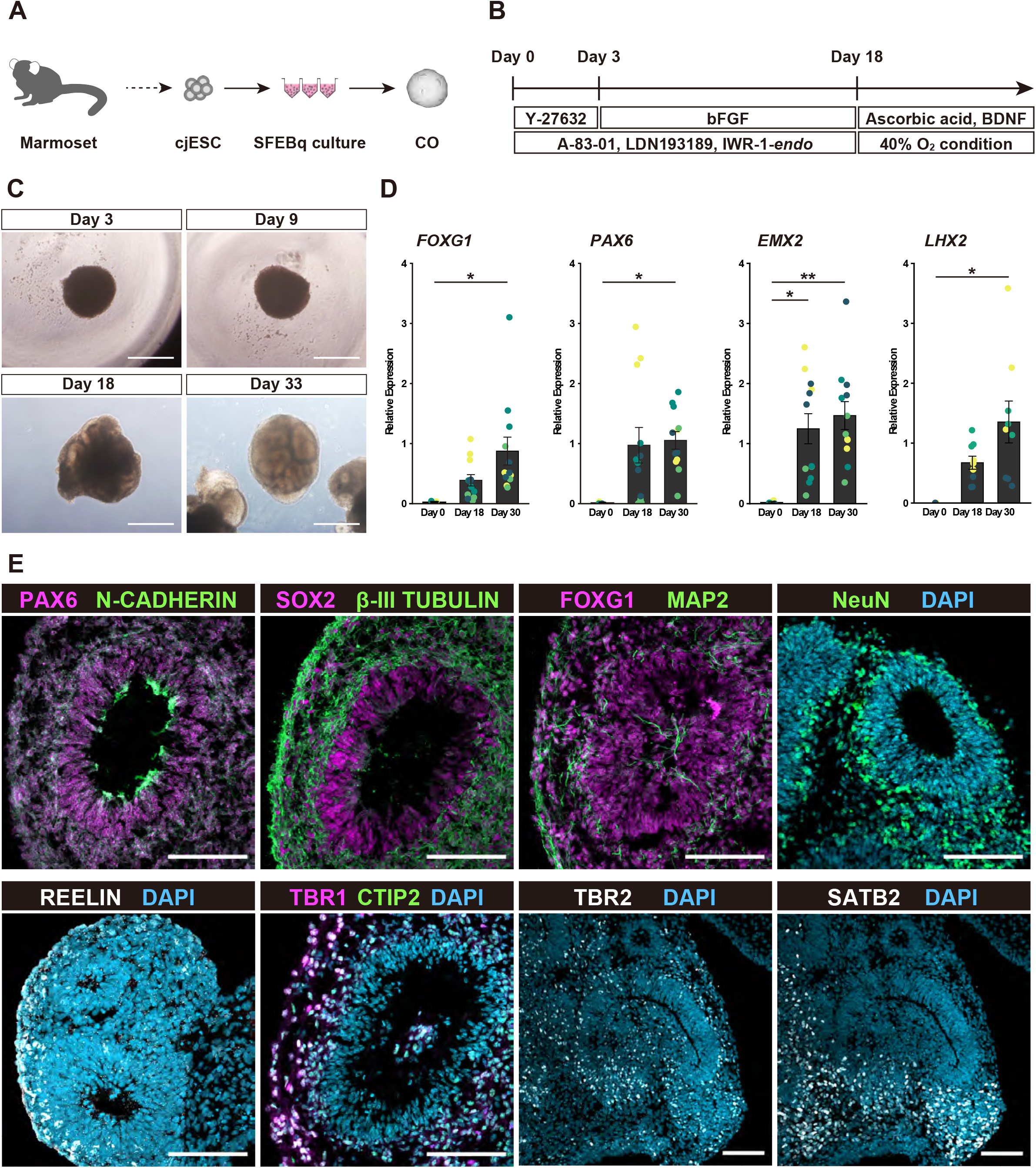
Cortical organoid generation and characterization. (A) CO generation from cjESCs. (B) A protocol to generate COs from cjESCs. (C) Bright-field images of cjESC-derived COs. Bars, 500 μm. (D) RT-qPCR for cortical markers. Each mRNA expression level was normalized to *GAPDH* expression (n = 9 or 12 organoids from three or four independent sets of experiments). \**P* < 0.05, \*\**P* < 0.01 vs. day 0. (E) Immunohistochemistry of COs on days 35, 47, and 64. Bars, 100 μm.

**Fig. 2:**
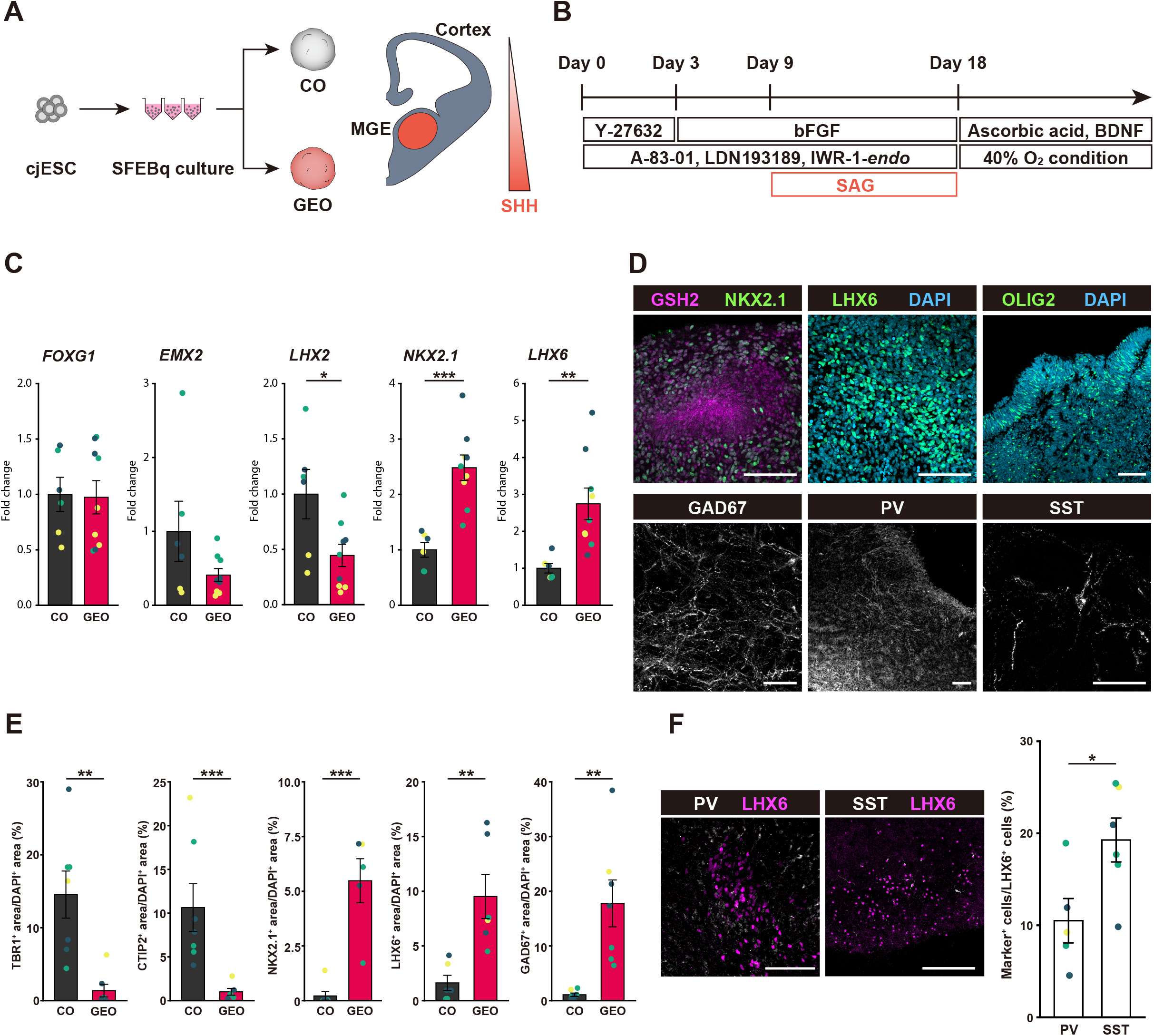
Ganglionic eminence organoid generation and characterization. (A) CO and GEO generation from cjES cells. (B) A protocol to generate GEOs from cjESCs. (C) RT-qPCR for dorsoventral markers. Each mRNA expression level was normalized to *GAPDH* expression. \**P* < 0.05, \*\**P* < 0.01 (n=6 or 9 organoids from three independent sets of experiments). (D) Immunohistochemistry of GEOs on days 47 and 64. Bars, 100 μm, 50 μm (GAD67 and SST). (E) Quantification of marker expression areas in immunohistochemistry on day 56. The area positive for each marker was normalized to the DAPI expression area. \*\**P* < 0.01, \*\*\**P* < 0.001 (n = 5–7 organoids from three independent sets of experiments). (F) Immunohistochemistry and quantification of PV^+^ or SST^+^ cells in the GEOs. \**P* < 0.05 (n = 5, 6 organoids from three independent sets of experiments). Bars, 100 μm (PV), 200 μm (SST).

**Fig. 3:**
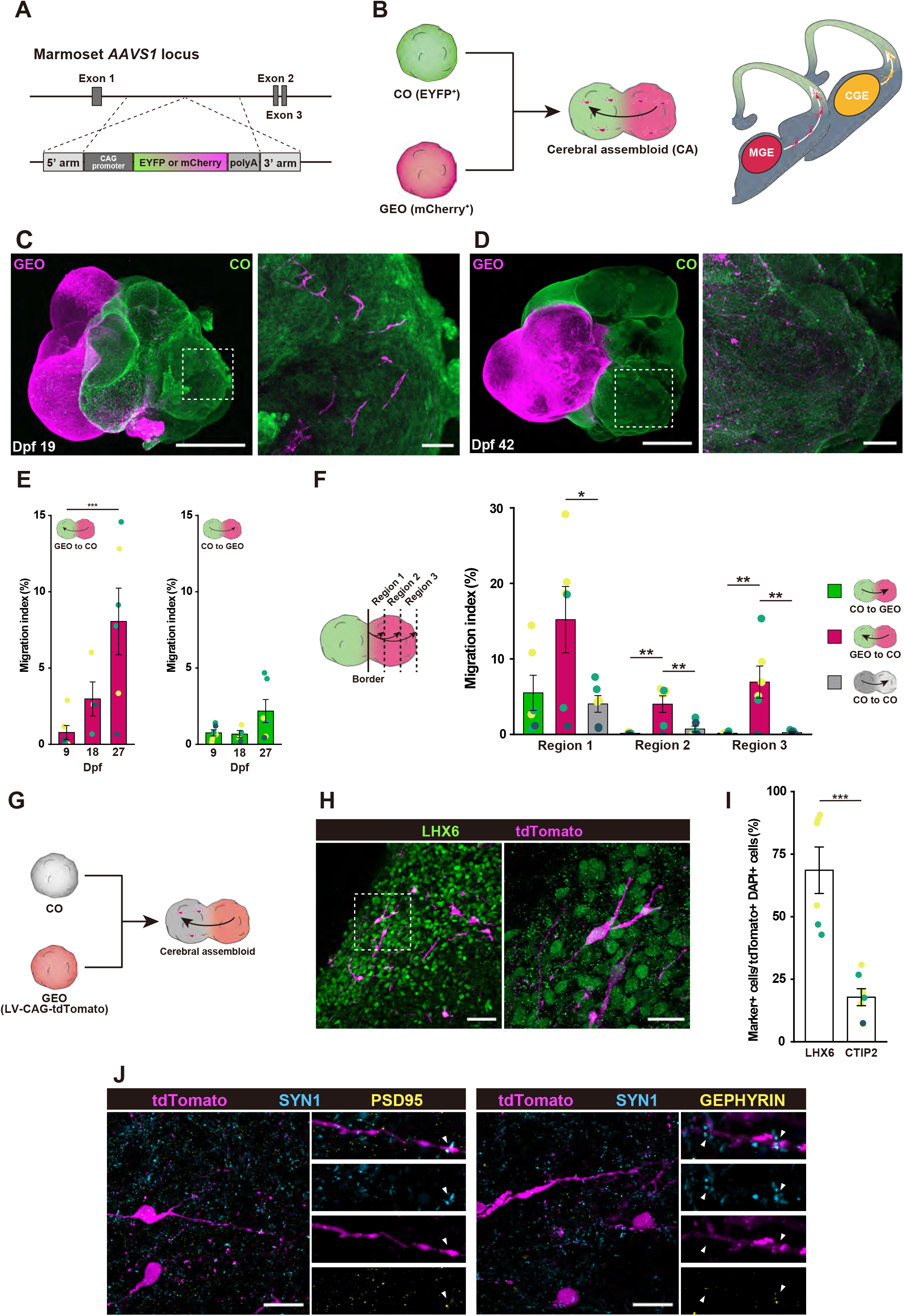
Cerebral assembloid generation and inhibitory neuron migration. (A) Genome editing at marmoset *AAVS1* locus to establish reporter cjESC lines. (B) CA generation and interneuron migration in the developing brain. (C, D) Whole immunofluorescent CA images. CUBIC-treated assembloids were stained at day 37 (19 days post-fusion; dpf) (C) and day 60 (42 dpf) (D). Right panels show magnification field of a white dotted line area in the left panel. Bars, 1000 μm (left panel), 100 μm (right panel in C), 200 μm (right panel in D). (E) Quantification of migration index in CAs at day 34, 43, and 52 (9, 18, and 27 dpf, respectively). The migration index was defined as the coverage proportion of migrated cell area in the contralateral organoid area. *** *P* < 0.001 (n = 4–6 CAs from three independent sets of experiments). (F) Quantification of migration index into counterpart region divided into three equal parts. \**P* < 0.05, \*\**P* < 0.01 (Tukey’s test, n = 6 CAs from three independent sets of experiments). (G) Cell migration assay in CAs. GEOs were virally labeled and then fused with nonlabeled COs to generate CAs. (H) Images of cortical side in virally labeled CAs at day 61 (30 dpf). tdTomato+ cells derived from GEOs expressed LHX6 in the CO side. Bars, 50 mm (left panel), 20 mm (right panel). (I) Quantification percentage of LHX6+ or CTIP2+ cells positive for tdTomato and DAPI (n = 6 CAs from two or three independent sets of experiments). (J) Immunohistochemistry of the cortical side in the virally labeled CAs at day 61 (30 dpf). Pre- and post-synapse markers (SYNI and PSD95/GEPHYRIN, respectively) were distributed along tdTomato+ neurites (white arrowheads). Bars, 20 μm.

## 3. Results

### 3.1. Generation of cortical and ganglionic eminence organoids from cjESCs

To generate COs from cjESCs, we optimized the SFEBq method [17] (Fig. 1A, S1A). The cjESCs were first dissociated into single cells and plated 10,000 cells to each well of a low-adhesion v-bottom plate. A higher Y-27632 concentration (50 μM) helped the robust formation of large, round aggregates of cjESCs (Fig. 1B, C). For efficient induction of the neuroectoderm and forebrain, the TGF-β signaling inhibitor A-83-01, the BMP signaling inhibitor LDN193189, and the Wnt pathway inhibitor IWR-1-*endo* were added from day 0–18. To characterize induced organoids, we performed qPCR analysis for cortical markers and found significant expressions of the early cortical marker *FOXG1* and the dorsal cortical makers *PAX6, EMX2,* and *LHX2* in the induced organoids (Fig. 1D), but little expressions of other germ layers, including the mesoderm and endoderm, and other brain regions (Fig. S1B, C). We also analyzed the organoid cytoarchitecture. Several round rosette clusters were found in the organoids. The neuroepithelium marker SOX2 was present in the inner zone of the rosettes on day 47, while cells positive for the neuronal markers βIII-TUBULIN, MAP2, and NeuN were distributed in the outer zone (Fig. 1E). The cortical marker FOXG1 was distributed across layers (Fig. 1E). The dorsal telencephalic progenitor marker PAX6 was also expressed in the inner zone. These results suggest polarized cortical neuroepithelial tissue formation in the induced organoids. During corticogenesis, layer-specific neurons are sequentially generated in the inside-out pattern in a time-dependent manner [34]. The expression of the very early-born neuron marker REELIN was found in the superficial layer on day 35 (Fig. 1E), suggesting the generation of Cajal-Retzius cells migrating to the superficial cortical zone. Layers VI and V neurons are then born from the cortical plate. The layer VI marker TBR1 and the layer V marker CTIP2 were expressed in the cell layers outside the ventricular zone on day 47 (Fig. 1E). At the later stages, upper-layer neurons are born. SATB2, an upper layer marker, was also observed in the outer zone (Fig. 1E). The subventricular zone (SVZ) is the hallmark of primates and contains radial glial cells and intermediate progenitor cells, which are responsible for the evolutionary neocortex expansion in neuron number and surface area [35]. TBR2, an intermediate progenitor cell marker, appeared in the induced organoids around day 60 (Fig. 1E). Few cells expressed the ventral telencephalic markers such as NKX2.1 or LHX6 (Fig. S1F). Organoids induced from cjESCs by our SFEBq-based protocol had typical dorsal cortical features, such as ventricular zone, cortical plate, and layer structure with deeper-layer and upper-layer neurons; hereafter, they are referred to as cortical organoids (COs).

Sonic hedgehog (SHH) signaling during telencephalic development is crucial for subpallial specification [36] (Fig. 2A). We therefore examined whether the dorsal-ventral specification of telencephalic progenitors in organoids could be controlled by SHH signaling activation during the SFEBq culture to induce the ganglionic eminence from cjESCs. SAG, a SHH signaling pathway agonist, was added from days 9–18 (Fig. 2B). SAG significantly increased the ventral telencephalic markers *NKX2.1* and *LHX6* expression in a concentration-dependent manner (Fig. S1D), whereas the dorsal telencephalic marker *EMX2* and *LHX2* expression decreased (Fig. 2C). We also assessed which ventral telencephalon regions were induced. We observed a few expression of other ventral telencephalon region (LGE, Striatum, CGE) markers (Fig. S1E). Immunostaining revealed that the SAG-treated organoids contained cells positive for the ventral telencephalic markers GSH2, NKX2.1, LHX6, and OLIG2, (Fig. 2D). These results suggest that SAG-treated forebrain organoids acquired ventral identity. Whether SAG-treated organoids further generated inhibitory neurons was also examined. On day 64, cells positive for the inhibitory-neuron-specific marker GAD67 and the inhibitory neuron subtype makers parvalbumin (PV) and somatostatin (SST) were present in these organoids (Fig. 2D). Additionally, more SST^+^ inhibitory neurons were generated in the organoids than PV^+^ inhibitory neuron (Fig. 2F). Only a few cells in the SAG-treated organoids expressed the deep layer neuron markers TBR1 or CTIP2 (Fig. 2E and S1G). Thus, the SAG-treated organoids have typical ganglionic eminence features and generate inhibitory neurons. Accordingly, SAG-treated organoids are referred to as GEOs.

### 3.2. Inhibitory neuron migration toward the cortex in cerebral assembloids

Inhibitory neurons generated from the GE undergo a long tangential migration to the dorsal cerebral cortex in the mammalian neonatal brain. To test whether this inter-regional interaction can be recapitulated *in vitro*, we generated assembloids with a dorsal-ventral axis by fusing COs and GEOs. Initially, two independent reporter lines of cjESCs were established using the CRISPR-Cas9 system to label each organoid with different colors. Fluorescent protein gene (EYFP or mCherry) was knocked in into the *AAVS1* locus, known as the safe harbor, which allows robust gene expression without exogenous gene silencing in almost all cell types (Fig. 3A, S2A-D). COs and GEOs were generated from EYFP-knocked-in cjESCs and mCherry-knocked-in cjESCs, respectively, and then fused on days 18–25 to produce cerebral assembloids (CAs) (Fig. 3B). mCherry-expressing cells were found inside the EYFP-expressing CO side 19 or 42 days after fusion, suggesting that GEO cells migrated toward COs (Fig. 3C, D). This cell migration was progressively increased and only in the GE to CO direction (Fig. 3E, S2E). These results suggest that the cell migration from GEOs to COs was unidirectional. We next determined how far cells migrated to the opposite region (Fig. 3F). Region 1 closest to the junction contained many neurons in every case, suggesting non-specific cell migration around the junction due to organoid fusion. CO cells were not distributed in regions 2 and 3 of the opposite GEOs. Notably, a significantly large number of cells derived from GEOs were found in regions 2 and 3 of the cortical side. Control experiment was also performed by generating cortico-cortical assembloids to determine whether this migration event is unique to the cortico-GE CA. A few CO-derived cells were observed in regions 2 and 3 of opposite COs in the cortico-cortical assembloids. Thus, the cell migration from the GE to the cortical side is unique to CAs. LHX6 is involved in migrating inhibitory neurons from the MGE to the cortex in the brain [37]. To verify that cell migration in CA mimics *in vivo* telencephalic development, whether migrating cells express LHX6 was examined. GEOs with tdTomato were first labeled by lentivirus expressing tdTomato under CAG promoter (VSV-G-LV-CAG-tdTomato) and then produced CAs by fusing lentivirus-labeled GEOs with unlabeled COs (Fig. 3G). tdTomato-expressing cells in the cortex side expressed LHX6 at 31 days after fusion (Fig. 3H, S2F), but not the cortical excitatory neuron marker CTIP2 (Fig. 3I, S2F). This migration inside the CAs could reflect GEO cells migrating to the cortex during *in vivo* telencephalic development. Furthermore, whether these migrating cells formed synapses in their CAs destination was examined. The pre-synapse marker SYNAPSIN I and either excitatory post-synapse marker PSD95 or inhibitory post-synapse marker GEPHYRIN were present across tdTomato^+^ neurites of GEO-derived cells in CAs on day 61 (Fig. 3J). These results suggest that only interneurons produced from GEOs can migrate to the COs and form synapses with cortical neurons in CAs. Thus, we conclude that the inter-regional interaction of the CAs along the dorsal-ventral axis recapitulates a long-distance interneuron migration from the ventral origin GE toward their target dorsal cortical region.

### 3.3. Developmental-stage-dependent neural activity patterns in COs and CAs

We next investigated whether neurons in the organoids generate neuronal activity. To obtain higher expressions of jGCaMP8m, a genetically encoded Ca^2+^ indicator, in CO neurons, tet-off system was used in two separate AAVs: AAV9-Esyn-tTA and AAV9-tetO-jGCaMP8m. tTA driven by neuron-specific synapsin promoter in mature neurons enhances the jGCaMP8m expression in organoids. Early-stage COs were infected with an AAV9-Esyn-tTA and AAV9-tetO-jGCaMP8m cocktail on day 47 or 48 and performed two-photon Ca^2+^ imaging 6 days after AAV infection. jGCaMP8m-expressing cells inside organoids spontaneously changed fluorescence signals (30–60 cells in 500 μm × 500 μm). These Ca^2+^ events were abolished by the Na^+^ channel blocker tetrodotoxin (data not shown). Fig. 4A represents typical Ca^2+^ dynamics traces in neurons of the early-stage COs (53-day organoids). The correlation analysis between all cell pairs in the field of view (Fig. 4B) and between frames throughout the time of the imaging (Fig. 4C, S3A, and Movie S1) revealed a high correlation coefficient between cells. These results indicate that many neurons at the early-stage COs exhibit synchronized neuronal activity. Two-photon Ca^2+^ imaging was also performed on late-stage COs (73–75-day organoids) to examine whether this synchronized activity persists or changes in mature organoids at a later stage. Late-stage organoids showed a high frequency of Ca^2+^ events (Fig. 4D). Interestingly, synchronized neural activity patterns were not observed. Correlations between all cell pairs and between frames during the imaging session were low (Fig. 4E, F, S3B, and Movie S2). The number of Ca^2+^ events were quantified and compared to assess whether neural activity patterns change as the organoids mature. The active neuron percentage at the later stage was lower than the early stage. Correlation analysis revealed that the correlation coefficient significantly decreased with organoid maturation (Fig. 4G-K). The sparseness of active neurons at the later stage was lower than at the early stage [38]. Principal component analysis was performed to evaluate neural activity pattern variability in these organoids. While the first principal component is much larger than other components in early-stage organoids, the first principal component explained only about 20% of the total variance in later-stage organoids (Fig. 4L). These results suggest that neurons in the late-stage organoids showed more mature, variable neural activity patterns than the early-stage organoids, similar to brain development [39,40]. Thus, COs can resemble developmental changes in neural activity patterns.

**Fig. 4:**
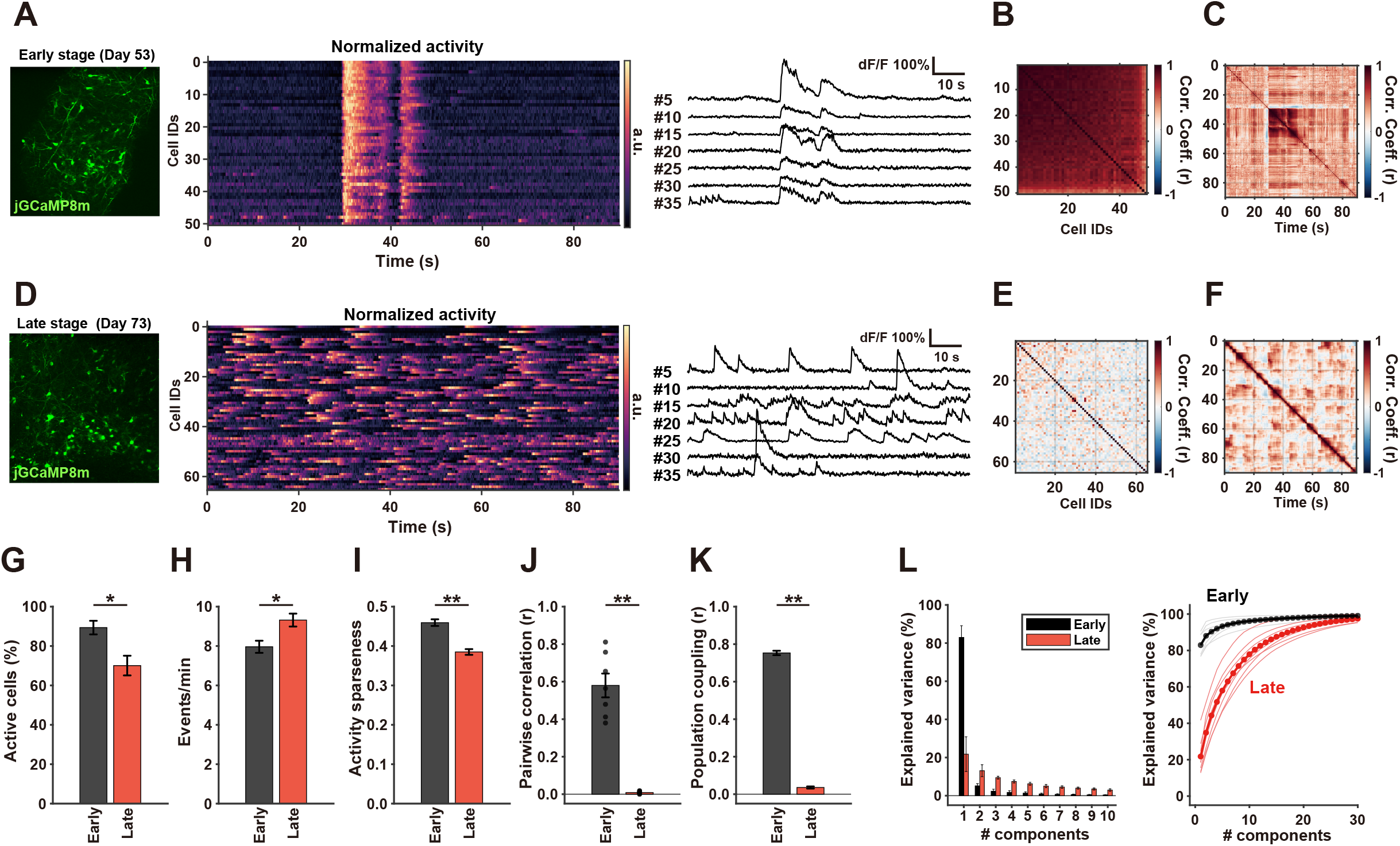
Developmental-stage-dependent changes in neural activity pattern in COs. (A) Neural activity in early-stage COs. Left: An image of a recording site. Middle: Normalized response matrix of all active neurons. Right: Time-series plots of ΔF/F in representative neurons. Cell order was sorted by population coupling values. (B) Correlation coefficient of activity between all pairs of simultaneously recorded neurons. (C) Correlation coefficient of activity patterns between all time-points during the recording. (D) Neural activity in late-stage COs. Left: An image of the recoding site. Middle: Normalized response matrix of all active neurons. Right: Time-series plots of ΔF/F in representative neurons. Cell order was sorted by population coupling values. (E) Correlation coefficient of activity patterns between all pairs of simultaneously recorded neurons. (F) Correlation coefficient of activity patterns between all time-points during the recording. (G) Active neuron ratio in early- and late-development stage organoids. (H) Ca event rate of individual neurons. (I) Activity sparseness of individual neurons. (J) Mean correlation coefficient statistics in each recording site. (K) Population coupling of individual neurons. (L) Principal component analysis of time-varied response patterns. Left: Percentages of explained variance for each principal component value of early-stage and late-stage COs. Right: Cumulative explained variance plot for each group (a solid bold or dotted line) and each imaging site (black or red thin line).

We next characterized the neural activity of the CAs. COs and GEOs were fused on days 30–33, applied the AAV9-Esyn-tTA and AAV9-tetO-jGCaMP8m cocktail on days 64–69, and performed Ca^2+^ imaging on the CAs on days 70–75. The cortex side of the CAs showed spontaneous unsynchronized activity (Fig. 5A-C, Movie S3). A significant difference was not found in neural activity patterns between late-stage COs and CAs except for the rate of Ca events during imaging (Fig. 5D-H). The decrease rate of Ca events in the CAs can be attributed to inhibitory inputs by migrated inhibitory neurons. The principal component analysis also demonstrated that CAs had an almost similar variance in activity patterns (Fig. 5I). Thus, CAs exhibited mature neural activity with an unsynchronized pattern. The late-stage COs and CAs showed similar unsynchronized activity patterns. This is presumably because the COs *per se* contain inhibitory neurons, as inferred from some LHX6 expressions in the COs (Fig. 2C, E) and the previous study based on single-cell RNA-seq [41]. It should be noteworthy that human cortical progenitors can produce both excitatory and inhibitory neurons, unlike mouse cortical progenitors [15,42]. Taken together, the marmoset CAs present a phenotypic neural network model of cortical dynamics.

**Fig. 5:**
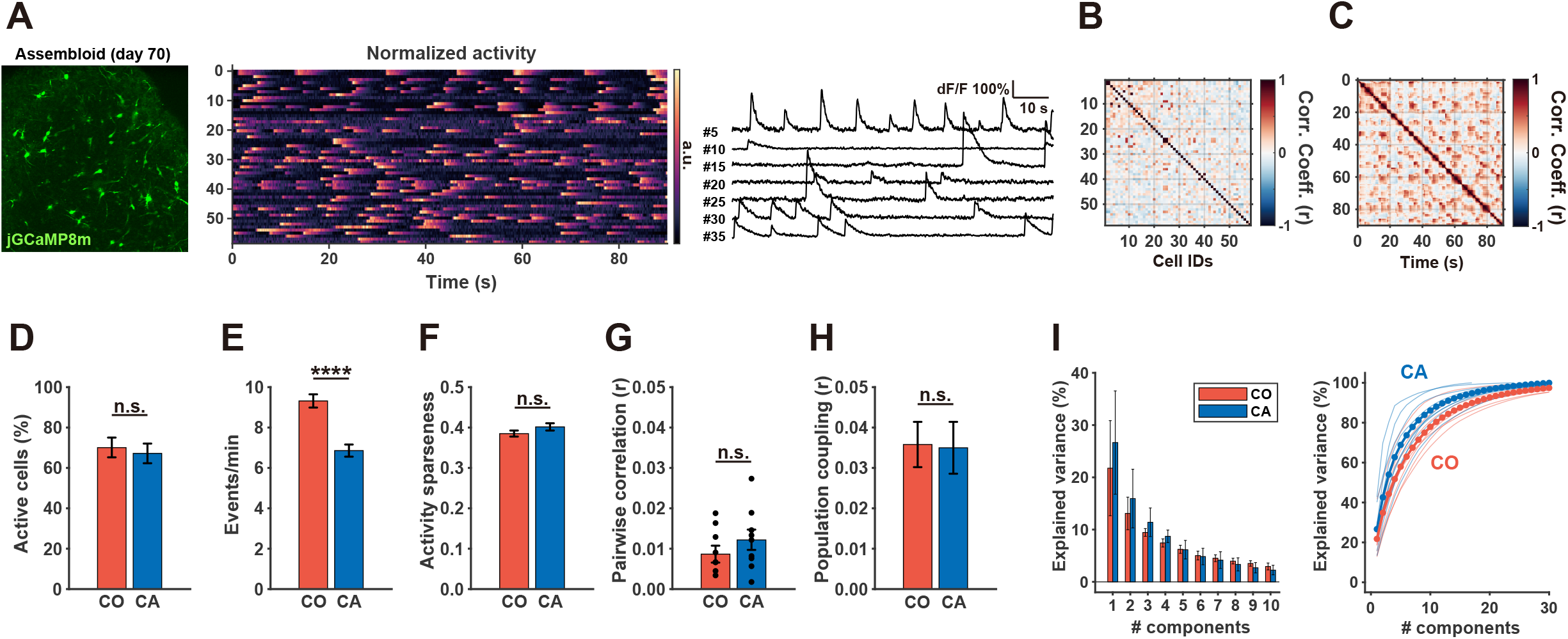
Unsynchronized spontaneous neural activity in CAs. (A) Neural activity in CAs. Left: An image of the recording site. Middle: Normalized response matrix of all active neurons. Right: Time-series plots of ΔF/F in representative neurons. Cell order was sorted by population coupling values. (B) Correlation coefficient of activity between all pairs of simultaneously recorded neurons. (C) Correlation coefficient of activity patterns between all time-points during the recording. (D) Active neuron ratio in late-stage COs and CAs. (E) Ca event rate of individual neurons. (F) Activity sparseness of individual neurons. (G) Mean correlation coefficient statistics in each recording site. (H) Population coupling of individual neurons. (I) Principal component analysis of time-varied response patterns. Left: Percentages of explained variance for each principal component value of late-stage COs and CAs. Right: cumulative explained variance plot for each group (Solid bold line and dots) and each imaging site (red or blue thin lines).

## 4. Discussion

Using an assembloid system derived from cjESCs, we established a marmoset brain model to recapitulate the long-distance migration of inhibitory neurons from the GE to the dorsal cortex and the cerebral cortex network activity. Cells derived from the GEO expressed LHX6, inhibitory neuron marker, and unidirectionally migrated toward the CO side in the CAs, while CO cells did not migrate to the GEO side. We also demonstrated changes in the CO neural activity with organoid maturation by two-photon Ca^2+^ imaging. At an early stage, COs showed synchronized burst activity. However, at late stage, COs exhibited asynchronized spontaneous activity with a reduced correlation between neurons. CAs showed mature neural activity with an unsynchronized pattern. Thus, we conclude that CAs derived from cjESCs resemble marmoset brain development in terms of the interaction between excitatory and inhibitory neurons.

CAs mimicked inhibitory neuron migration during development. GE organoid-derived LHX6+ cells began to migrate into the cortical side in the CAs at day 37 (19 days after fusion) when the marginal and ventricular zone (VZ) formed in the CO (Fig. 1E). During mouse brain development, GE-derived inhibitory neurons enter the cortex around embryonic day 12.5 (E12.5)[43], when TBR1^+^ deep layer neurons appear between marginal zone and VZ. Comparing this mouse corticogenesis period with our marmoset organoid system, marmoset organoids day 37 seems to correspond to E12.5 mouse brain. Marmoset CAs will allow us to study inhibitory neuron migration mechanisms *in vitro*.

COs and CAs recapitulated the cerebral cortex network activity. The changes in neural activity during CO maturation reflected developmental-stage-dependent neural activity *in vivo* [39]. Notably, CAs exhibited mature unsynchronized activity patterns. The neural activity pattern changes during brain development and under pathological conditions. At the early developmental stage, synchronized burst activities transiently appear by electrical coupling via gap junctions. This synchronous activity is considered to aid in proliferation, migration, and axon guidance. Normal mature brain exhibits asynchronized spontaneous activity by chemical synapses mediated by NMDA, AMPA, or kainate receptors [44]. These unsynchronized activity patterns at later stages are crucial for neural circuit maturation. The timing and mechanisms of development-dependent activity change have been reported in mice, but not in marmosets [40]. In contrast, the epileptic state during a seizure shows synchronized activity as well as an imbalance between excitation and inhibition (E/I) in the cerebral cortex. E/I imbalance is considered a common disease mechanism for autism spectrum disorders and epilepsy [45]. Marmoset CAs can be an *in vitro* model complementary to *in vivo* physiology. How activity changes shape neural development, circuit organization, and pathogenesis of neurological and psychiatric disorders remains unresolved. The three-dimensional microphysiological systems using CAs should provide mechanistic insight into this question.

A next challenge in CAs will be cell-type-specific circuit organization. Inhibitory neurons in the cerebral cortex are diverse and classified into three sub-cell types with biochemical markers: PV, SST, and vasoactive intestinal protein (VIP) [12]. Each cortical interneuron sub-cell type has distinct synaptic targeting biases onto neighboring excitatory pyramidal neurons and engages in its own circuit motif. In the cerebral cortex, PV^+^ cells form inhibitory synapses on the perisomatic regions of pyramidal neurons, control the action potential generation and timing in pyramidal neurons, and subserve feedforward inhibition [46]. SST^+^ cells synaptically connect with pyramidal neuron distal dendrites and selectively inhibit specific inputs to pyramidal neurons, mediating feedback inhibition [46]. VIP^+^ neurons target SST^+^ inhibitory neurons and mediate inhibition of inhibition, so-called disinhibition, on pyramidal neurons. The formation of connectivity and circuit motifs of cortical local circuits in the assembloids will be warranted for further investigations.

The assembloid system derived from cjESCs can be applied to different brain regions by taking advantage of ESC ability to differentiate into diverse cell types corresponding to different brain region identities. The marmoset assembloid system using ESCs and iPSCs will provide a platform for elucidating molecular, cellular, and circuit mechanisms of the NHP brain and also bridge rodent research and translation into humans in neuroscience research, regenerative medicine, and drug discovery.

## Acknowledgments

We would like to thank the members of Osakada Laboratory for their stimulating discussions. This study was supported by Grants-in-Aid from the Japan Society for the Promotion of Science (F.O., K.S.), Brain/MINDS from the Japan Agency for Medical Research and Development (F.O., A.A.: JP15dm0207032), and CREST from the Japan Science and Technology Agency (F.O.).

## Author contributions

T.K. cultured cjESCs, generated organoids and assembloids, performed genome editing, virus production, and imaging experiments, collected and analyzed all data, and wrote the manuscript. R.F.T. analyzed imaging data. S.T. cultured cjESC lines. K.S. assisted with genome editing. H.K. and A.A. contributed to the knock-in strategy at the AAVS1 locus. S.S. and H.O. contributed to cjESC culture. F.O. wrote the manuscript and supervised the project.

---

Supplemental Movie 1

Two-photon calcium imaging on the early-stage cortical organoids

Supplemental Movie 2

Two-photon calcium imaging on the late-stage cortical organoids

Supplemental Movie 3

Two-photon calcium imaging on the cerebral assembloids on day 70

## Supplementary Figures

**Fig. S1.**
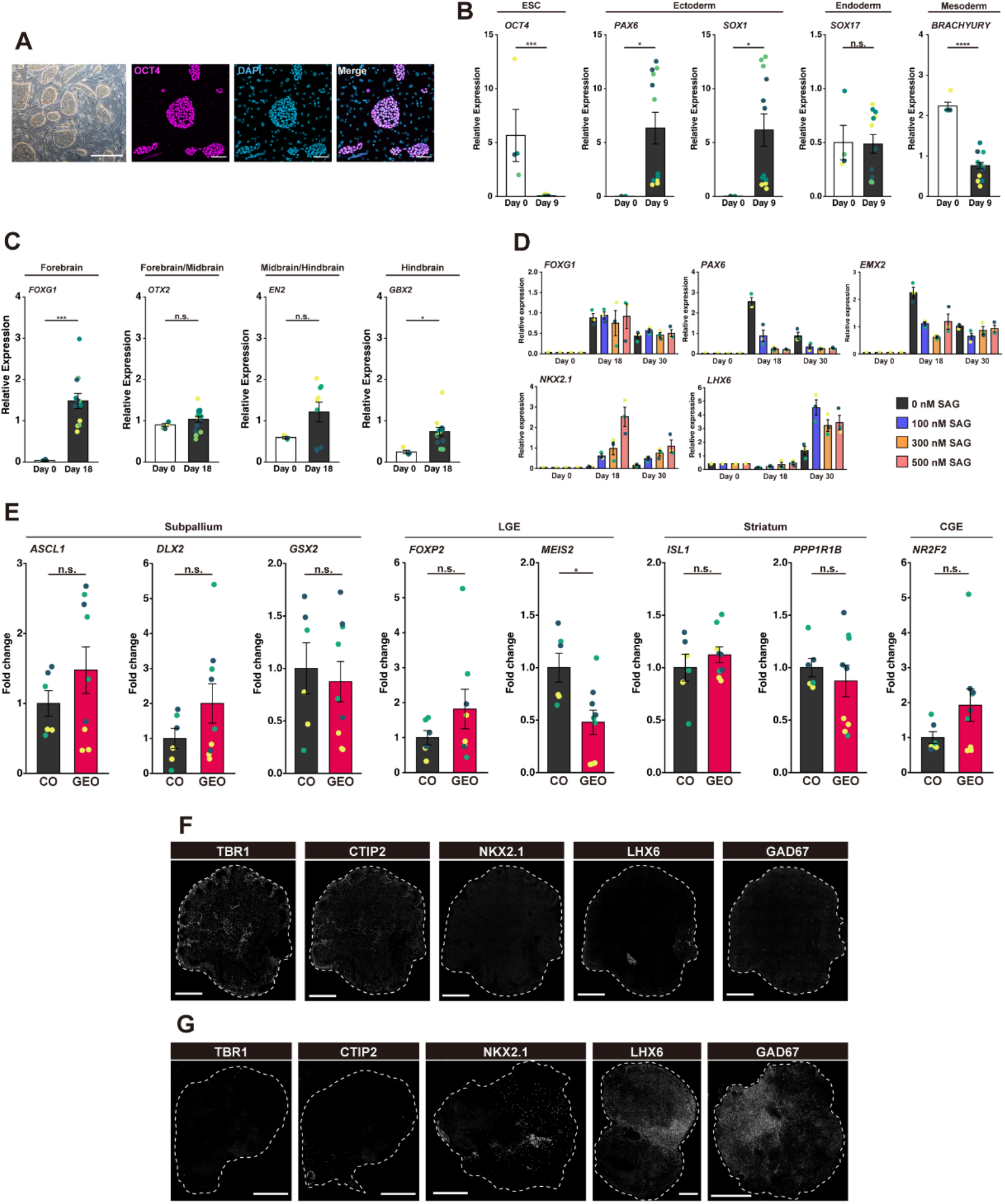
Characterization of cortical organoids and ganglionic eminence organoids, related to Figures 1 and 2. (A) A bright-field image and immunofluorescent images of undifferentiated cjESCs. Scale bars, 500 μm (Bright field) and 100 μm (Immunostaining). (B) Quantification of mRNA expression by RT-qPCR on day 9. Each mRNA expression level was normalized to *GAPDH* expression. \**P* < 0.05, \*\*\**P* < 0.001 (*t*-test, for *FOXG1, OTX2, GBX2:* n = 4 (Day 0) or n = 12 (Day 18) from four independent sets of experiments, for *EN2:* n = 3 (Day 0) or n = 9 (Day 18) from three independent sets of experiments). (C) Quantification of mRNA expression by RT-qPCR on day 18. Each mRNA expression level was normalized to *GAPDH* expression. \**P* < 0.05, \*\*\**P* < 0.001, \*\*\*\**P* < 0.0001). (D) Quantification of mRNA expression by RT-qPCR. Each mRNA expression level was normalized to *GAPDH* expression (n = 3). (E) Quantification of mRNA expression by RT-qPCR. Each mRNA expression level was normalized to *GAPDH* expression. \**P* < 0.05 (*t*-test, n = 6 or 9 from three independent sets of experiments). (F, G) Immunohistochemistry of COs (F) and GEOs (G) on day 56. Each representative image was used for quantification analysis in Fig. 2E. Scale bars, 500 μm.

**Fig. S2.**
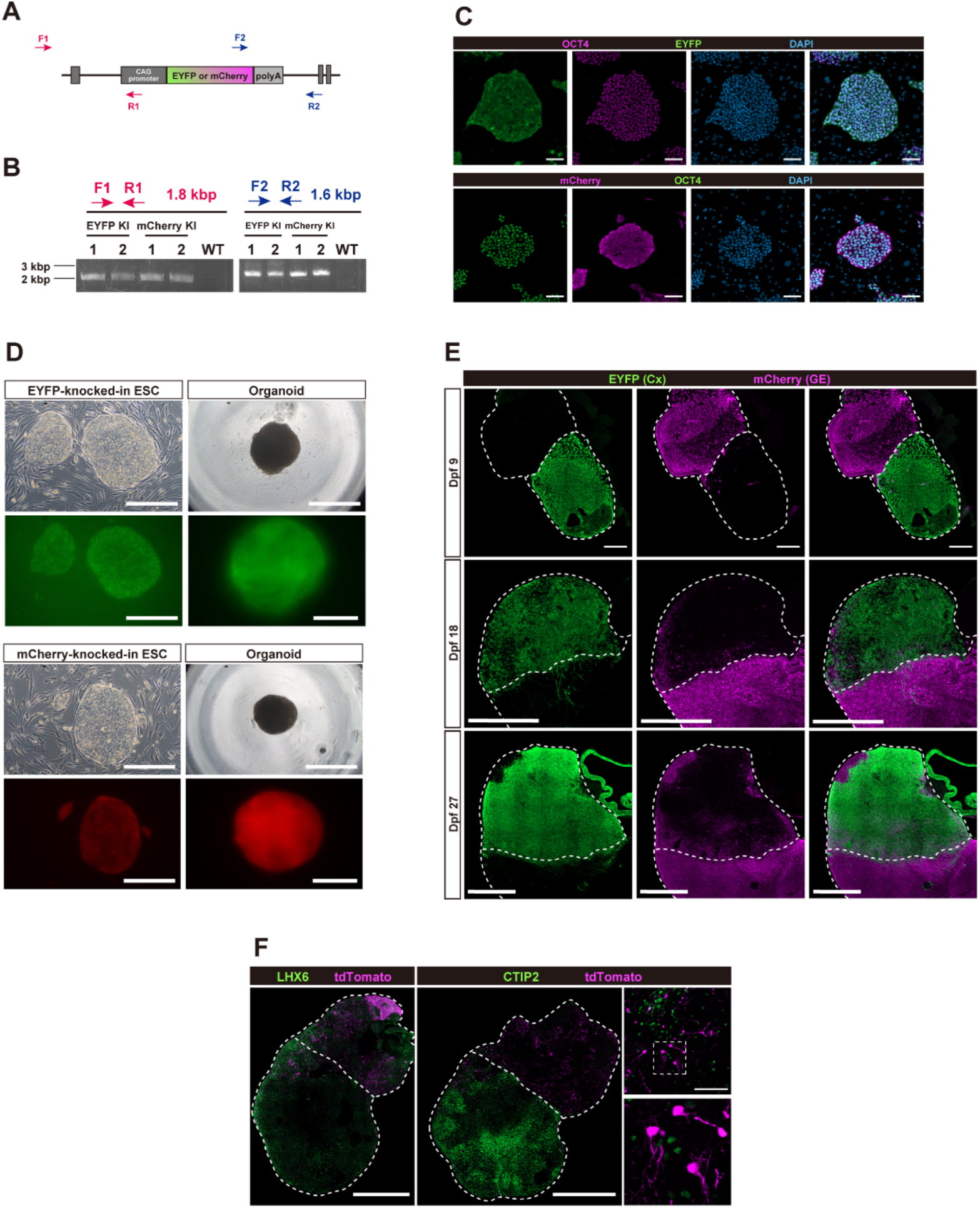
Migration analysis in cerebral assembloids, related to Fig. 3. (A) A schematic diagram of sequence primers for detection of knock-in. Arrows show the primer sets for genotyping. (B) Genotyping of the knocked-in cjESCs by PCR. These primer sets do not anneal with WT genomic locus. (C) Immunofluorescent images of knocked-in cjESC lines. Knocked-in cjESCs expressed OCT4, the pluripotent marker, and a fluorescent protein (EYFP or mCherry). Scale bars, 50 μm. (D) Bright-field images and fluorescent images of knocked-in cjESCs and organoids. Scale bars, 500 μm, 200 μm (organoid fluorescent images). (E) Representative images of cerebral assembloids at each time point. Scale bars, 200 μm (dpf 9), 500 μm (dpf 18), and 1000 μm (dpf 27). (F) Representative images of immunohistochemistry of virally labeled cerebral assembloids at day 61 (dpf 30). White dotted line shows outline and border of assembloids. Scale bars, 500 μm and 50 μm.

**Fig. S3.**
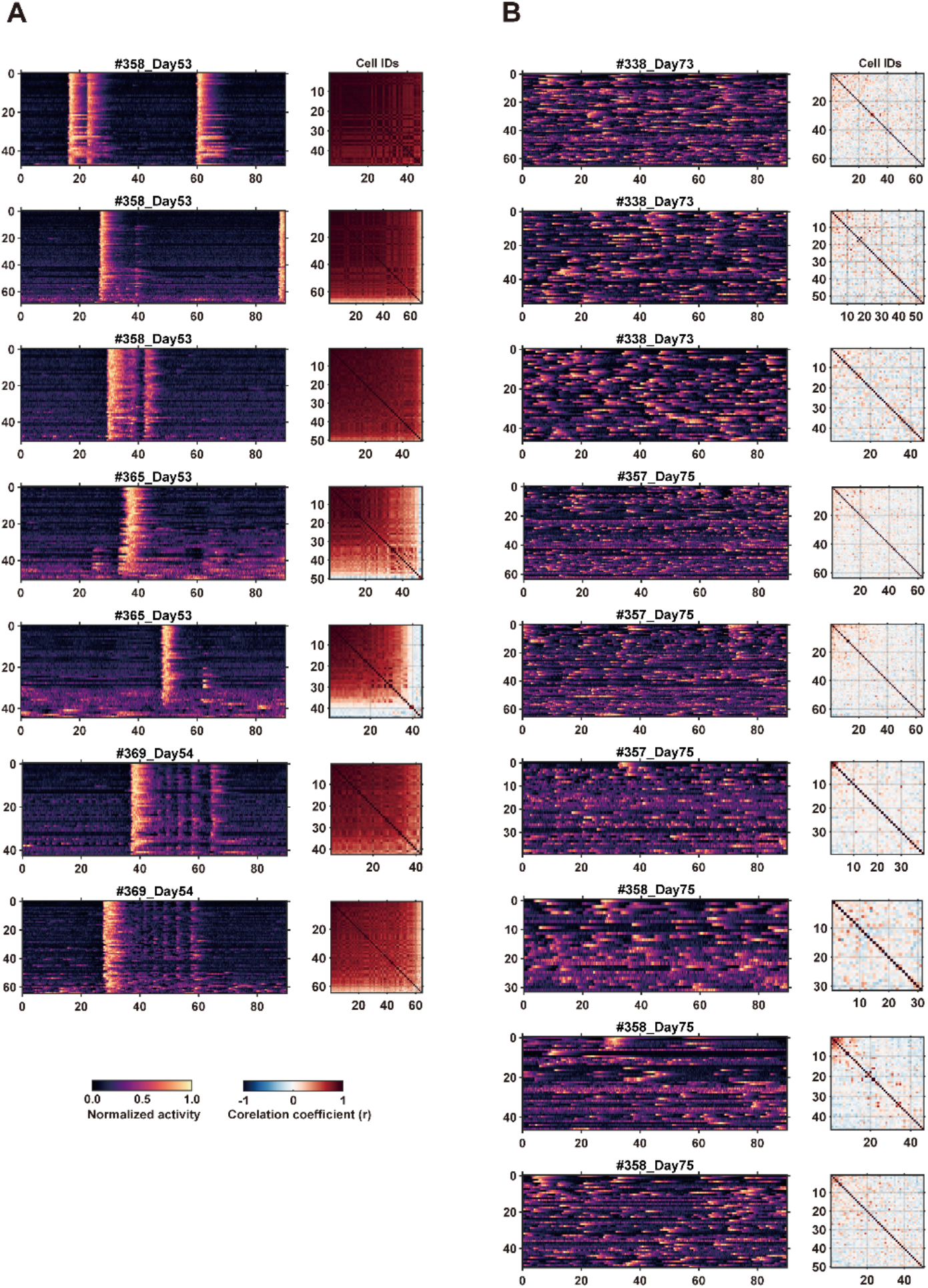
Calcium imaging of early- or late-stage cortical organoids, related to Fig. 4. (A, B) Normalized response matrix of all active neurons in each recording site (left), and correlation coefficient of activity between all pairs of simultaneously recorded neurons (right) in early-stage (A) or late-stage (B) COs.

**Supplementary Table1.**
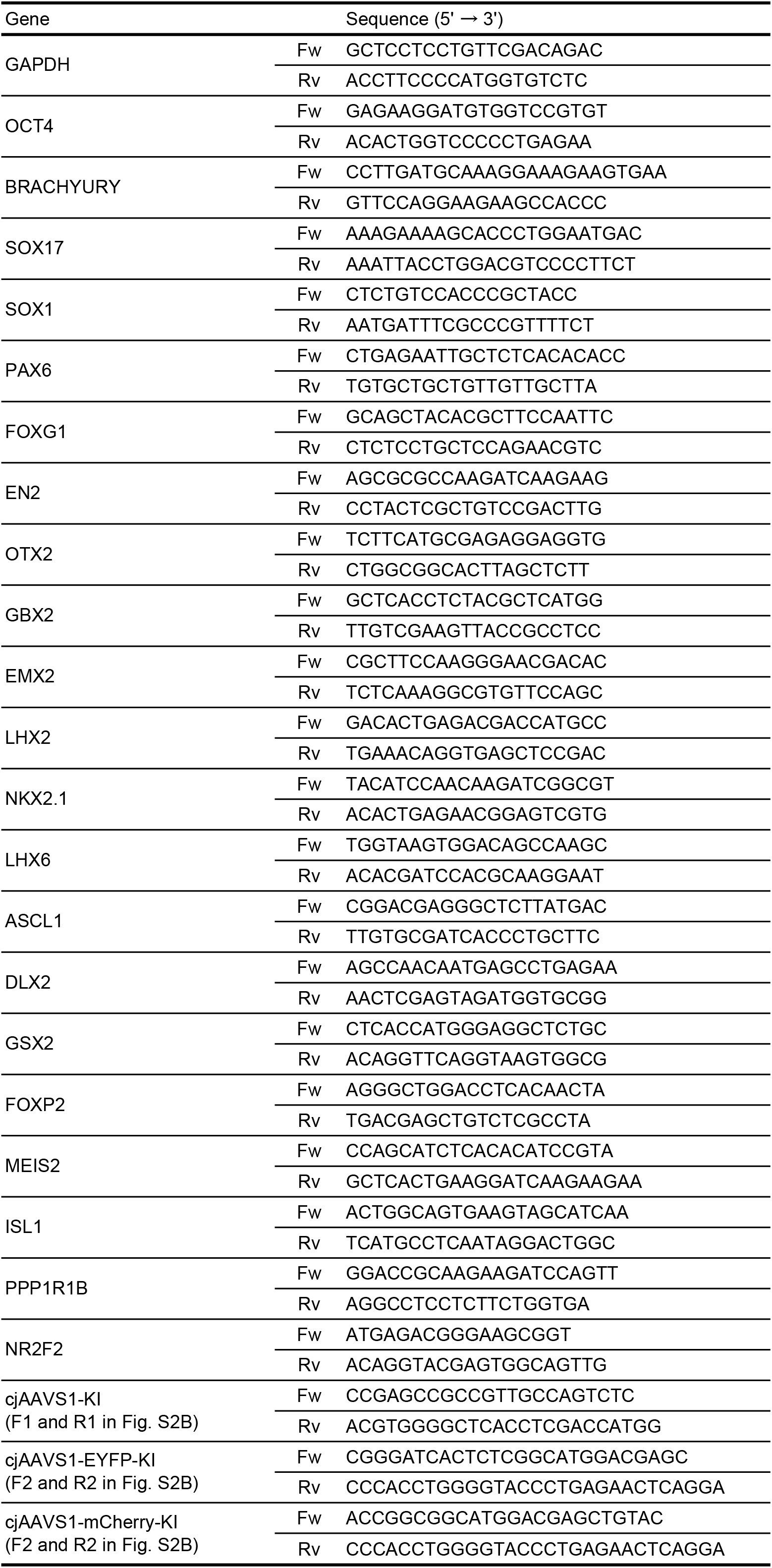
Primers for qPCR analysis

